# Dynamical persistence in resource-consumer models

**DOI:** 10.1101/2020.02.18.954008

**Authors:** Itay Dalmedigos, Guy Bunin

## Abstract

We show how highly-diverse ecological communities may display persistent abundance fluctuations, when interacting through resource competition and subjected to migration from a species pool. This turns out to be closely related to the ratio of realized species diversity to the number of resources. This ratio is set by competition, through the balance between species being pushed out and invading. When this ratio is smaller than one, dynamics will reach stable equilibria. When this ratio is larger than one, fixed-points are either unstable or marginally stable, as expected by the competitive exclusion principle. If they are unstable, the system is repelled from fixed points, and abundances forever fluctuate. While marginally-stable fixed points are in principle allowed and predicted by some models, they become structurally unstable at high diversity. This means that even small changes to the model, such as non-linearities in how resources combine to generate species’ growth, will result in persistent abundance fluctuations.

## I. INTRODUCTION

Resource competition is one of the main mechanisms underlying species interactions. Theoretical works [1, 2] have demonstrated that communities interacting via resource competition may exhibit different dynamical behaviors (also observed in nature [3, 4]), including relaxation to equilibria, limit cycles and chaotic dynamics. In systems consisting of a few species and resources, the dynamical outcome may depend on all the details describing the interactions in the community [5]. For systems with higher dimensionality (more species and resources), full detailed knowledge of the interactions may be difficult to obtain and predictions might seem hopeless, and potentially sensitive to all unknown details.

In recent years, research on high-dimensional communities has shown that full knowledge on all the interactions might not always be needed [6–14], and important ecological quantities such as total biomass and diversity can be predicted from a handful of statistics on the interaction parameters. In the space of these relevant statistics, one can identify different regions (known as “phases”) with qualitatively distinct behaviors, such as relaxation to equilibria versus chaotic dynamics. Within these phases, the qualitative behavior is robust, i.e. insensitive to sufficiently small changes in the systems’ interaction coefficients.

In this paper, we consider high-dimensional communities with resource-competition interactions. We show that in an entire region of parameter space, the system fails to reach equilibria and instead abundances fluctuate indefinitely. This might seem surprising, as some theoretical models are known to always lead to stable equilibria, including classical models by MacArthur [15, 16]. We argue that when the number of species and resources is large, there are regions of parameter space where these models are highly sensitive (structurally unstable), and even very small changes to the model will result in persistent abundance fluctuations.

A key ingredient in our discussion is competitive exclusion, according to which the number of species that can coexist in a stable equilibrium is smaller or equal to the number of resources (or more generally, the number of niches). A *marginally*-stable fixed point *can* accommodate more species than resources, but it can be destroyed by small perturbations or changes to the dynamical rules.

The sensitivity of marginally-stable equilibria raises the following question: what then replaces the marginally stable fixed-point, once it is no longer stable? There are two possible scenarios: (1) Species will go extinct until an equilibrium with fewer species is reached, which satisfies the competitive exclusion principle, or The system will not reach any fixed point, and instead abundances will continue to fluctuate indefinitely. We show that for a community experiencing migration from a species pool, the generic situation is number (2) above.

Resource competition dynamics in diverse communities have been analyzed in a number of works employing tools from statistical physics [7, 10, 13, 17]. For a region of model parameters, marginally-stable [10, 18] or close to marginally-stable [7, 13] equilibria are reached. Yet the models studied all admit a unique equilibrium by construction (in the spirit of classical works [15, 16]). For example, species’ growth rates are assumed to depend linearly on resource availability, which cannot accommodate effects such as essential resources [19]. A model combining resource-competition with other interactions not mediated by resources, was studied in [7]. It showed that a unique stable equilibrium cannot exist in a certain region of parameter space, but did not study what replaces that unique equilibrium. Additional factors might drive communities to marginal stability, such as metabolic trade-offs [20] or evolution, highlighting the importance of studying the generic dynamics in these situations.

Our argument proceeds as follows. Interactions create a balance between species being pushed out due to competition, and species invading when they can, steering the community towards some target species richness. If this richness is larger than the number of resources, then fixed points generically will be unstable, and persist abundance fluctuations will ensue, See Fig. 1(C). These dynamics are characterized by species being pushed out by fixed points’ instability, and back when they are able to invade.

**Figure 1.**
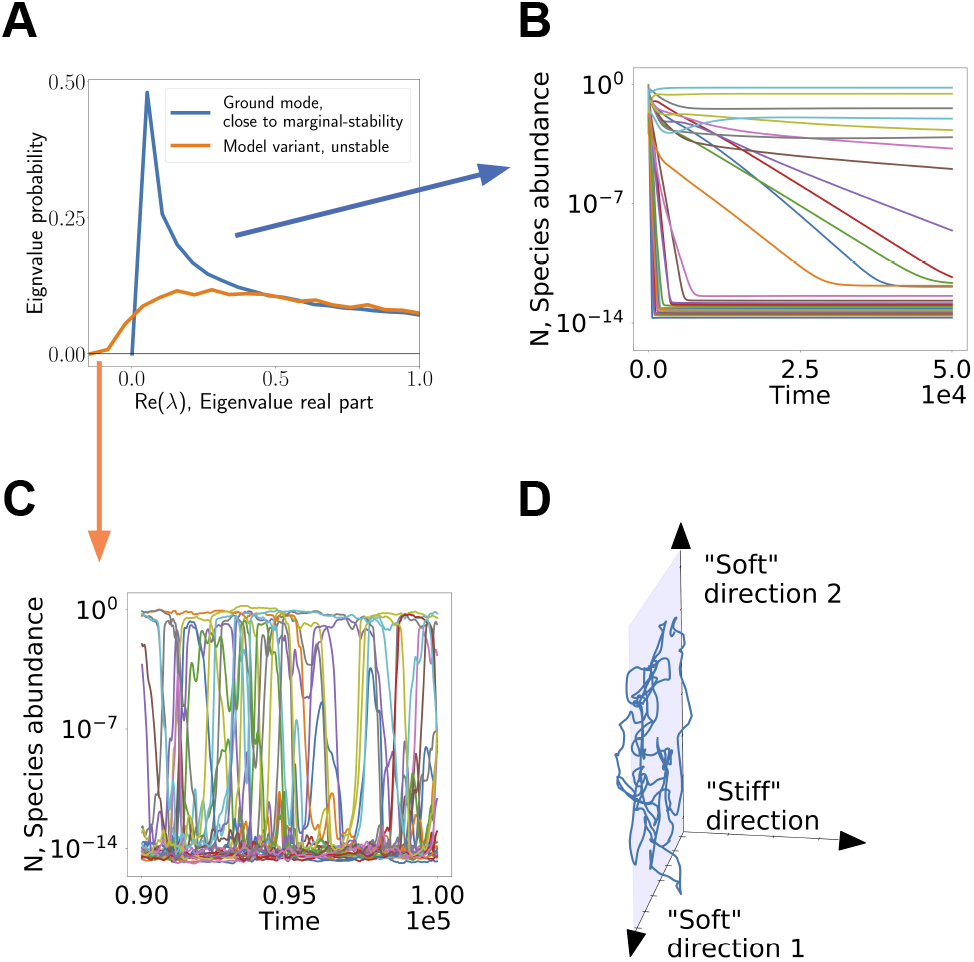
Summary of argument. (A) The fixed points encountered by a high-diversity resource-competition community may be unstable. In the presence of migration this creates persistent abundance fluctuations, shown in (C), in which species are pushed out due to the instability, but are later able to invade again. (B) Marginally-stable fixed points (or nearly marginal ones) that appear in certain models, are characterized by a non-negative spectrum, also shown in (A). They allow the community to relax to a fixed point. But the stability of such fixed points is sensitive to modeling assumptions, including additional interactions of other types, or how growth rates depend on resource availability. Introducing these will generically push the system towards unstable non-equilibrium dynamics shown in (C). (D) In such a case, the directions corresponding to the nearly marginal eigenvectors become “soft” directions, showing large fluctuations. For clarity, in (B,C) 30 representative species are plotted.

This instability is manifested by the spectrum of response to small perturbations around a putative fixed point at the target species richness, Fig. 1(A). Under certain modeling assumptions, these fixed points might be marginally stable, but in this case small changes to the model push the fixed point to become truly unstable, without changing much the target richness set by the competition, see Fig. 1(C). In other words, it is precisely the large number of (nearly-)marginal directions that allows for such fluctuating dynamics to persist, as shown Fig. 1(D). Marginal, or nearly-marginal eigenvectors around the fixed point become “soft” directions, namely combined abundance fluctuations of multiple species that are met with little resistance. This correspondence is further explored in Appendix F.

The paper is structured as follows. Sec. II A defines the ground model used to illustrate the arguments. Sec. II B looks at the effect of changes to the model, by adding interactions on top of resource competition. It shows how the dynamics generated by this model might vary significantly due to even small changes, replacing equilibria by non-equilibrium dynamics. The general mechanism behind this sensitivity is explained in Sec. II C. In Sec. II D, the behavior is shown to be sensitive in a second variant of the model in which all interactions are strictly the result of resource competition, but with non-linear resource intake.

Sec. II E describes the resulting abundance distributions and community diversity. The non-equilibrium coexistence of more species than there are resources or niches, is of great interest in its own right. It has been suggested to play a part in the resolution of the “paradox of the plankton” [21]. In Sec. II E we consider this question directly in a high-dimensional setting, in light of works on high-dimensional chaos in well-mixed communities [6, 14] and meta-communities [22, 23]. Finally, Sec. III concludes with a discussion, focusing on predictions for experiments and natural communities.

## II. METHODS AND RESULTS

### A. The ground model

To illustrate the ideas we use a well-known model and introduce two variants to that model. The canonical model is MacArthur’s resource consumer model (MCRM) [15], that will be referred below as the “ground model”. The variants introduce small changes to its dynamical evolution.

The MCRM describes the dynamics of *S* species abundances *N*_*i*_ (*i* = 1 … *S*) competing over *M* types of resources *R*_*β*_ (*β* = 1 … *M*). The MCRM system evolves according the following set of coupled differential equations

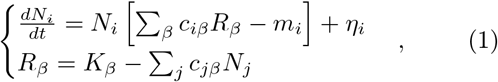

where *c*_*iβ*_ describes the consumption preference of species *i* for resource *β*. *m*_*i*_ is a minimum maintenance cost that must be met by species *i* for it to grow. *K*_*β*_ is the carrying capacity of resource *β*. The first equation includes a migration term *η*_*i*_ from a species pool. It will taken to be small, allowing species to invade if they have positive growth rates. Plugging the expression for *R*_*β*_ into the first equation yields

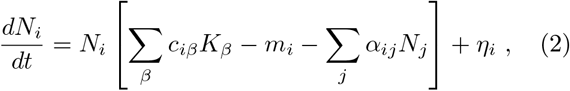

where 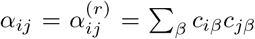, and the superscript (*r*) denotes resource-mediated interactions. This equation is now in the form of generalized Lotka-Voltera equations.

### B. Sensitivity to direct interactions: a demonstration

A key result by MacArthur [15] is that the model in Eq. (2) exhibits globally stable dynamics, reaching a single fixed point independently of the system’s initial conditions. In this section we show that by a small addition of other interactions on top of the resource competition interactions described above, the system is no longer guaranteed to approach a fixed point. Instead, for a broad region of control parameters, the species’ abundances fluctuate indefinitely, see Fig. 1(C).

To demonstrate this phenomena we introduce the first variant of the MCRM, which includes additional “direct” species interactions, 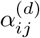, so that the total interaction coefficients read 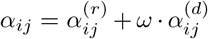, with *ω* controlling the strength of the perturbation. These direct interactions may come as a result of many mechanisms that lie beyond the unperturbed MCRM. The important point will be to find when such additional interactions have a large effect on the dynamics, even when they are small.

To quantify the size of the perturbation, we use the ratio of the Frobenius norms (sum of squared interaction coefficients) of the interaction matrices, setting ‖*ω* × *α*^(*d*)^‖ _*F*_ / ‖*α*^(*r*)^‖ _*F*_ = 0.05 throughout. For any given model parameters, *ω* is chosen satisfy this condition, allowing for a comparison between results with different model parameters.

The quantities *c*_*iβ*_, *K*_*β*_, *m*_*i*_, 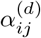 that define the interactions are drawn at random, representing a generic diverse community, without any additional structure beyond that already incorporated into the resource-competition model. The parameters *c*_*iβ*_, *m*_*i*_, *K*_*β*_ are drawn independently for each value, and 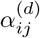 are drawn independently, except possibly a correlation between 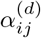 and 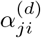 controlling the symmetry of the direct interactions. All quantities are drawn from Gaussian distributions, parameterized by their first two moments. The definitions of parameters are given in Appendix A.

As seen in Fig. 1(C), the variant with even these small additional interactions shows persistent abundance fluctuations, even if the ground model reaches equilibrium, Fig. 1(B).

### C. Theory for the onset of non-equilibrium dynamics

To understand whether and when the variant of the model will reach a fixed point or a non-equilibrium state, we look at the stability of fixed points, assuming they are reached. The basic idea is that systems are sensitive to perturbations, if the fixed point in the ground model is close to marginal stability.

Before turning to the present model, we review the results for the random Lotka-Volterra models, which in the terminology of Sec. II B only have “direct” interactions, 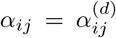. Their dynamics have been studied recently [8, 9, 12, 14] (see also related results in other models [6]).

A number of sharply delineated regions in parameter space are found, referred to below as ‘phases’. In one phase the system reaches a unique equilibrium. The boundary of this phase is marked by loss of stability of these fixed points. Beyond this boundary (with a sharp transition at large *S*) lies another phase, where the dynamics fail to reach a fixed point and abundances fluctuate indefinitely [14]. In a special case where the interactions are symmetric, namely *α*_*ij*_ = *α*_*ji*_, this phase is instead characterized by with many possible alternative equilibria, all of which are close to marginal stability [12].

The behavior of the model variants defined above and in Sec. II D, bares many similarities to that of the random Lotka-Volterra models. There is a unique equilibrium phase, which is delineated by a boundary at which the equilibrium looses its stability. Beyond it, we find in simulations that the dynamics never reach a fixed point, as shown in Figs. 1,4. The case of symmetric 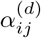 is special and appears to follow the scenario in [12], see Appendix H. An important difference from random Lotka-Volterra models is the mechanism by which fixed points loose their stability, which we now discuss.

We describe a method of calculating the species richness and stability of the fixed points for the model variant described in Sec. II B. This method is exact when the system admits a unique fixed point; the loss of its stability marks the boundary of the phase. To highlight the relation between marginal stability and sensitivity to perturbations, we study the spectrum of the interaction matrix, and how it changes for the model variant described above. A different approach, using Dynamical Mean Field Theory, is possible and ultimately equivalent, and has been employed on a related problem in [7].

Consider fixed points of the dynamics, i.e. abundance vectors 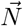 for which 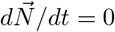 in Eq. (2). We are interested in the linear stability of these fixed points, namely whether the system approaches the fixed point if initialized close to it. The linear stability can be obtained from the properties of the reduced interaction matrix *α*\* comprised only from interactions between surviving species (for which *N*_*i*_ → *c* > 0, even as the migration *η*_*i*_ → 0). A fixed point 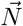 is linearly stable if and only if all the real parts of eigenvalues of *α*\* are positive, or equivalently if the minimal eigenvalue real part is positive, 0 < min {Re [Λ (*α*\*)]} ≡ *λ*_*min*_. A fixed point is marginally stable if *λ*_*min*_ → 0^+^ when *S* → ∞.

While the MCRM only has stable or marginally stable fixed points, *λ*_*min*_ ≥ 0, the model variant can have unstable ones. Close to marginality, i.e. when *λ*_*min*_ is zero or close to zero, even a small perturbation may cause the system to lose its stability. This is the case for a broad region in parameter space, as we now show.

In Fig. 2, we show the spectrum of such an *α*\* matrix close to a marginal fixed point. As expected, the marginal case is characterized by non-vanishing density of eigenvalues arbitrarily close to zero. When applying a small perturbation to the marginal interaction matrix *α*\*, for example 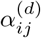 described in Sec. II B, the spectrum is broadened and may cross zero to give eigenvalues with negative real parts, resulting in a dynamically unstable fixed point. The properties of the fixed point depend crucially on the species richness (the number of species that survive), which is a result of a balance between competition that pushes species out of the system, and migration which allow them to try and invade.

**Figure 2.**
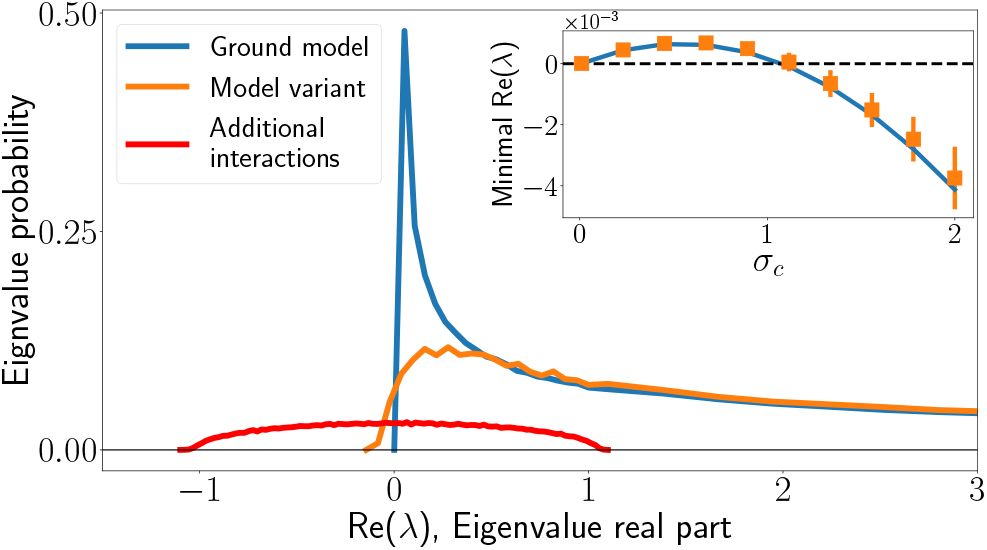
Spectrum of *α*\*, the interaction matrix of persistent species, in the ground model (blue), and the variant with additional direct interactions (orange). The perturbation spectrum is shown in red, to illustrate its size we normalize the area under the perturbation spectrum to the size of the relative perturbation strength (0.05). (Inset) Minimal eigenvalue real part of the reduced interaction matrix *α*\*, when varying *σ*_*c*_ at fixed *μ*_*c*_. Solid line is theoretical curve. A phase transition occurs when the minimal eigenvalue real part crosses from *λ*_*min*_ > 0 at which fixed points of the dynamical system are stable, to *λ*_*min*_ < 0 where all fixed points of the system are unstable, leading to persistent dynamics.

The method for calculating the spectrum consists of the following main steps: first, we find the number *S*\* of coexisting species using the cavity method. This follows similar calculations in precious works [9, 13], and is detailed in Appendix C. We then calculate *λ*_*min*_, the minimal real part of the eigenvalues of *α*\*, for a reduced interaction matrix with *S*\* species. This is done using random matrix theory and detailed in Appendix D.

Following this method, we can predict the dynamical behavior as a function of the model parameters. We find three phases, shown in Fig. 3. In the first, the system converges to a unique fixed point, independent of the initial conditions, as in Fig. 1(B). In the second, the system fails to reach a fixed point, with abundances fluctuating indefinitely, as in Fig. 1(C). In the third phase the abundances diverge, indicating that the model is no longer adequate in this parameter regime.

**Figure 3.**
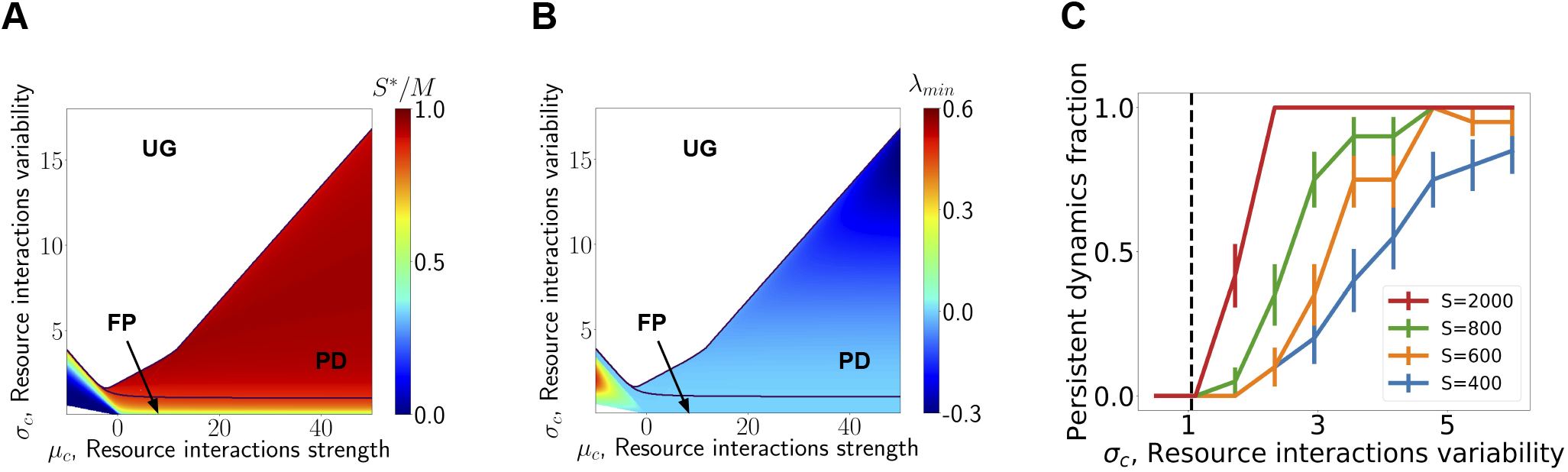
The model exhibits three phases, i.e. regions with qualitatively distinct behavior. In one, the system converges to a stable fixed point (FP), in another fixed points of the system are unstable yielding persistent dynamics (PD). In the third phase, unbounded growth (UG), species abundances grow without bound. *ω* is adjusted in order to maintain constant perturbation strength of 0.05. (A) Color map of the ratio *S*\**/M*, indicating how close the system is to competitive exclusion *S*\**/M* = 1. (B) The minimal real part of eigenvalues of the interaction matrix between coexisting species, *λ*_*min*_. Fixed point stability is lost at *λ*_*min*_ = 0, resulting in a phase transition to dynamically persistent states. (C) Probability for having a persistent dynamics in a system with interactions drawn for different values of *σ*_*c*_, the variability in consumer preferences. The transition between FP and PD phases becomes sharper as system size increases.

Notably, when *S*\**/M* ≈ 1 the unperturbed system is close to competitive exclusion and correspondingly close to marginality, and therefore the model variant with the direct interactions becomes unstable, i.e. *λ*_*min*_ < 0. As expected theoretically, the transition between the two behaviors is sharp when *S, M* are large, and happens at the theoretically predicted value of the parameters, see Fig. 3(C). In less diverse systems, the transition is more gradual.

The loss of stability of putative fixed points results in persistent dynamics where species invade but are then pushed back out by the instability of fixed points. This is clear in Fig. 1(C).

### D. Resource-competition with non-linear resource intake

So far, we have discussed the ground model with a small addition of other interactions. This allows us to identify regions in parameter space where the ground model is sensitive to perturbations. By adding interactions that are not mediated by resource competition, the model variant can no longer be strictly interpreted as a resource-competition model. Here we consider a second variant of the ground model, which belongs to the resource-competition class, but with non-linear resource intake. Non-linear dependence of the growth rate on resource availability appears in many situations, such as in competition over essential resources [19, 24]. Here the aim is not to study the consequences of a specific non-linear mechanism, but rather to demonstrate the sensitivity of the model to such variations in the dynamical rules.

We find that much like the model variant discussed in previous sections, here too the dynamics are sensitive to the changes from the ground model, with fixed points turning into persistent abundance fluctuations, in much the same parameter regions as found previously.

The second variant to the ground model, Eq. (1), is different from the ground model in the way that different resources translate into the growth rate of the consumer. Whereas in Eq. (1) the growth rate is a linear combination of the resource values *R*_*β*_, here we use a non-linear function. We choose a non-linear consumption function 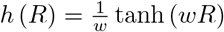 with control parameter *w*. With this consumption function, the dynamical equations read:

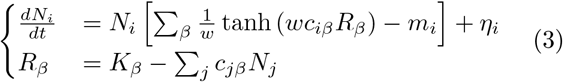

The parameter *w* allows us to tune the deviation from the ground model. For small values of *w*, *h* (*R*) ≃ *R* so the non-linear effects become small and the equations reduce to the ground model Eq. (1). For finite *w* non-linear effects may be important. We quantify the deviation from linearity explored by the dynamics by 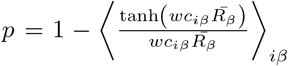, where (*‥*)_*iβ*_ is the average over species and resources, and 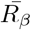 denotes the time average of the resource abundance *R*_*β*_ (taken over a window of Δ*t* = 1000). We find that even a rather small value of *p* is sufficient to induce a transition to dynamical persistence, see Fig. 4, where the transition occurs at *p* ~ 0.06. Again, this demonstrates how the system’s dynamics may be sensitive to small changes in the equations governing the model, in this case in a variant that is itself strictly a resource competition model.

**Figure 4.**
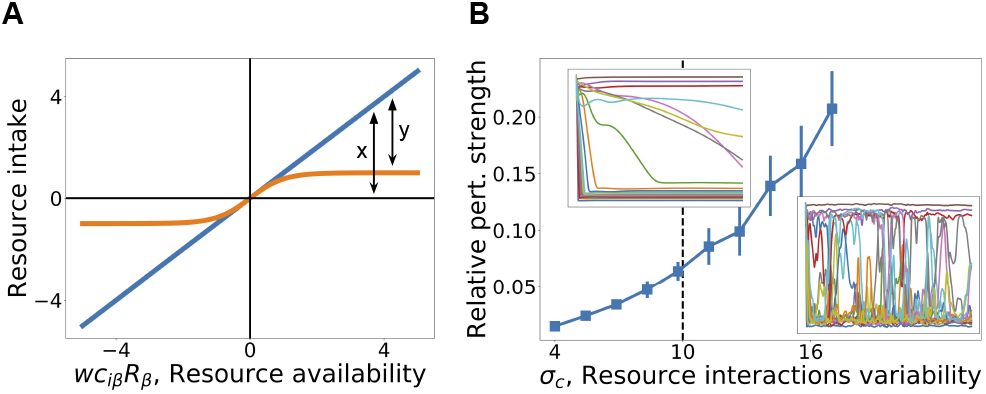
A model with non-linear resource intake. (A) Illustration of the non-linearity. The weighted linear sum of the intake *wc*_*iβ*_*R*_*β*_ of resource *β* by species *i*, used in the ground model, is replaced by a non-linear function. The level of non-linearity, *p*, is measured by the ratio *y/x*, see figure, where the *x* axis denotes the intake and the *y*-axis the linear- and non-linear intake functions, averaged over species and time. (B) *p* changes as model parameters are changed (here varying *σc*). Persistent non-equilibrium dynamics are found for *p* ≳ 0.06.

### E. Species abundance distribution and diversity

Above, we saw how systems near marginal stability are sensitive to small variations in the model, either by additional interactions, or by changes to the functional form of the interactions. Here we show that these changes can allow the diversity to go well above the number of resources. This is made possible by the persistent dynamics, which are no longer bound by the competitive exclusion principle.

The competitive exclusion principle [25] states that for models describing an ecological community of *S* species relying on *M* limiting resources, no stable fixed points with *M < S*\* exist. Briefly, the core of the argument is that any fixed point with *M < S*\* would imply a degenerate Jacobian matrix with rank *M* or less. This kind of fixed point can be marginally stable, but not stable. The second variant of the model, Eq. (3), satisfies the conditions for this principle to hold, so the diversity of stable equilibria is bound by *M*.

As an example we look at the second model variant, as defined in Sec. II D. Long-time simulations of the persistent dynamics show that the species abundance distribution converges to a stationary form that can be decomposed into a power law at intermediate abundances, and other parts at the highest and lowest abundances, see Fig. 5(A):

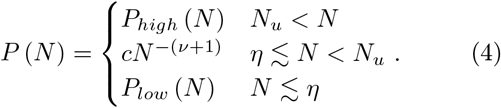

**Figure 5.**
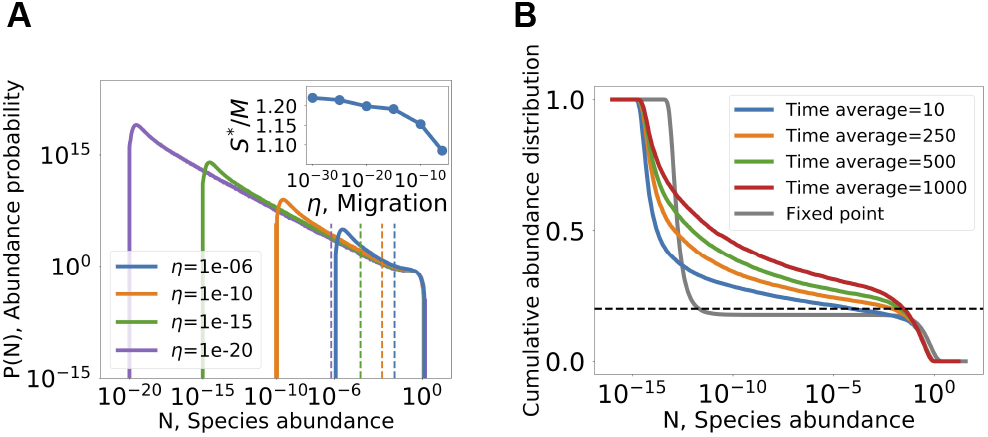
Diversity above competitive exclusion in the non-linear RC model II D. (A) Abundance distribution for different values of migration. The area to the right of the vertical lines hold exactly *M* species. The rest of the distribution, below the line, accounts for species above competitive exclusion. Inset: the total number of coexisting species normalized by number of resources, with values above one indicate crossing of competitive exclusion. (B) Due to the abundance fluctuations, averaging abundances over a time window pushes the distribution of abundance upwards due to fluctuations. The cumulative abundance distribution is shown, defined as 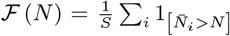. The dashed line is the competitive exclusion bound, *M/S*. For comparison, the distribution at a fixed point is given, showing that the number of species at high abundance (*N* ≳ 10^−3^) does not reach this bound, and the rest of the species are at low abundances, only supported by migration.

Here *N*_*u*_ is a constant, and *c* is set by the normalization ∫*P* (*N*) *dN* = 1. From the simulations, *ν* is not far from zero when *η* → 0 (*ν* ≈ 0.02 in Fig. 5, and similar for other parameter sets, see Appendix B).

We first ask about the instantaneous species richness, namely the fraction of species that are not at the migration floor (say, above 100*η*). By integrating *P* (*N*) in Eq. (4) one finds that the fraction of the species above the migration floor approaches a finite number when *η* → 0^+^, for details see Appendix E. This number can be larger than *M*, and in fact is so in the example shown in Fig. 5. In other words, a finite fraction of the species coexist above the competitive exclusion limit even when migration is very small. This is possible since the community is not in a fixed point, and so is not bound by the competitive exclusion principle.

If the species abundance is measured by integrating over a finite-time window, see Fig. 5, the abundances shift to higher values as the time window grows, indicating that species have periods of time with high abundance. This leads to a growth in the abundance *N*_*CE*_ above which there are exactly *M* species with higher abundances.

In [23], chaotic dynamics where studied in Lotka-Volterra equations with random interactions coefficients, and the existence of of time periods with high abundance have been reported, as well as a power law like in Eq. (4) (albeit with a different exponent). The relation of these results to the present resource-competition model are an interesting question for future research.

## III. DISCUSSION AND CONCLUSIONS

In this work we have shown how diverse ecological communities with resource-competition interactions may display non-equilibrium dynamics. This turns out to be closely related to the ratio of realized species diversity to the number of resources, *S*\**/M*. When this number is larger than one, fixed-points are either unstable or marginally stable, as expected by the competitive exclusion principle. If they are unstable, the system is pushed away from fixed p oints, a nd a bundances forever fluctuate. W hile m arginal-stable fi xed po ints are in principle possible, they are structurally unstable under variations in the model, such as non-linearities that destabilize the fixed points.

### Comparison with random Lotka-Volterra models

This picture bridges a gap to the behavior of high-dimensional models where interactions are sampled at random without a specified mechanism. In the notation of Sec. II B, this corresponds to having 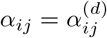 only. These models show a phase with persistent dynamics [6–9, 12, 14], in contrast to resource-competition which have thus far only shown relaxation to equilibrium in highly diverse communities [10, 13]. We find that the generic phase-diagram is in fact much more similar, with a transition to non-equilibrium dynamics when the variability in interaction strengths is high enough (compare, for example, Fig. 3 with the phase diagram in [9]).

One difference is that here, the symmetry of the interactions can be very high and still lead to non-equilibrium dynamics. For example, in the model with added direct interactions (Sec. II B), the total interactions are very close to symmetric, with corr (*α*_*ij*_, *α*_*ji*_) = 0.997. In random Lotka-Volterra models, dynamics at a comparable level of symmetry would typically relax to equilibria. This highlights the importance of certain structures in the interaction network on dynamics.

### Predictions

How can the behavior discussed in this work be identified in natural or experimental communities? The dynamical outcome will depend on the following considerations:

- Is the community isolated; under migration from a regional species pool; or part of a meta-community?
- The ratio of realized species diversity to the number of resources (*S*\**/M*).
- Is the realized diversity *S*\* high enough for high-diversity effects to show?

Consider first a single well-mixed system with continuous migration from a species pool, which was the focus of previous sections. In such a setting, dynamics either a relax to single uninvadable equilibria or reach persistent fluctuations. Which of these two possibilities is realized depends on the system parameters: the realized species diversity (*S*\*) is set by the balance between extinctions due to competition and species able to invade. If fixed points at this diversity are unstable, the latter outcome will result. This is the Persistent Dynamics phase in Fig. 3. As shown in Fig. 3, it is attained when there is sufficient variability in the interactions, mediated for example by a broad distribution of consumption preferences (high *σ*_*c*_).

We turn to a single well-mixed community that is isolated (no migration, *η*_*i*_ = 0). Here species may go extinct due to large abundance fluctuations, without being able to invade again. Extinctions may then lead to equilibria even when non-equilibrium dynamics are expected with migration, see Appendix G. The difference is that these equilibria can be invaded by species from the species pool. Importantly, in these conditions *all* fixed points are invadable, as uninvadable ones would translate to equilibria in the presence of migration. If there are now isolated migration events from the species pool that are well-separated in time (for example, at low migration rates, or in experiments where species are re-introduced) the equilibria will be punctuated by migration events that change the community composition [26].

An explicit spatial dimension, such as a meta-community in which several well-mixed systems are coupled by migration, again changes the phenomenology. In this case, one might also find persistent fluctuations for a meta-community, even if it is isolated from any outside species pool, allowing species to go extinct within it. Still, the remaining species might continue to fluctuate for extremely long times without inducing extinctions. This has been shown recently for many-species meta-communities with random Lotka-Volterra interactions in [22, 23]. An example simulation, provided as a proof-of-principle, is provided in Appendix G. The conditions for non-equilibrium dynamics to persist depend on additional parameters including the migration rates and the number of communities in the meta-community. A fuller account of this effect is an interesting direction for future research.

Finally, we note that the non-equilibrium dynamics discussed in this work apply to communities with many species and resources or niches. Simulations indicate that dynamical fluctuations appear when there are tens of species in the community or more; communities with fewer species may instead relax to equilibria.

High-dimensional ecological dynamics are, in some respects, qualitatively different from their low-dimensional counterparts. Here we classified possible scenarios for the dynamics of resource-competition communities, and provided predictions for each scenario. We hope it may help in guiding future theoretical works, observations and experiments on high-diversity communities.

## Acknowledgments

It is a pleasure to thank J.-F. Arnoldi, M. Barbier and G. Biroli for helpful discussions. G. Bunin acknowledges support by the Israel Science Foundation (ISF) Grant no. 773/18.

## Appendix A Basic setup

Each instance of the first model variant, defined in Sec. II B, requires setting the values of the quantities 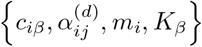, referred to here as the disorder parameters. They are fixed from the control parameters {*S, M, μ*_*c*_, *σ*_*c*_, *μ*_*d*_, *σ*_*d*_, *γ*, *μ*_*m*_, *σ*_*m*_, *μ*_*K*_, *σ*_*K*_} as follows.

Let us denote by 〈*X*〉 the expectation value of random variable *X*. All results cited in the paper at high-diversity depend only on the first and second moment of the system disorder parameters distribution. The means and variances of these are given by 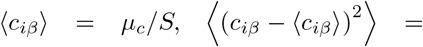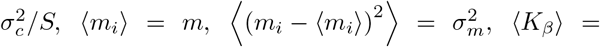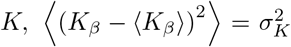, and the direct interactions by 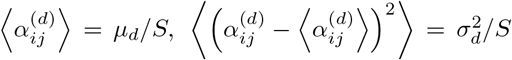 and 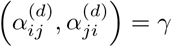 −1 ≤ *γ* ≤ 1. All other cumulants are set to zero. This definition of the parameters ensures that the abundance distribution *P*(*N*_*i*_) and the fraction of persistent species have a finite, well-defined limit as *S, M* are taken to be large. In other words, in that limit all results will only depend on these control parameter combinations, e.g. on *μ*_*c*_ = *S* 〈*c*_*iβ*_〉 rather than on *S, 〈c*_*iβ*_ separately. The same results are obtained for a sparse interaction matrix with *C* non-zero links per species, as long as 1 ≪ *C*. In that case, which includes the case *C* = *S* above, the moments are rescaled by *C* rather than *S*, e.g. 〈*c*_*iβ*_〉 = *μ*_*c*_/*C* instead of (*c*_*iβ*_) = *μ*_*c*_/*S*.

To simplify the notation below, it is useful to separate quantities into mean and fluctuating parts

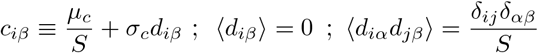

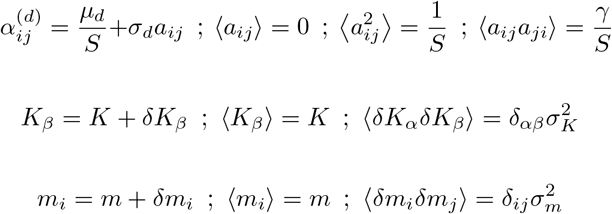

With these definitions the first variant, Eq. (2), can be

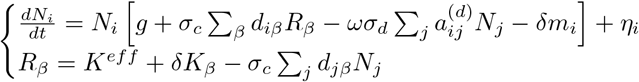

where

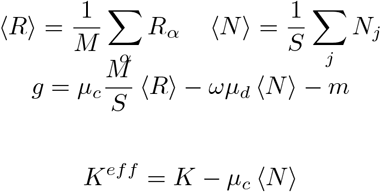

This form of the equations will be useful below, in Appendix C.

## Appendix B Model definition, parameters and simulation details

Differential equations were integrated using a Radau integrator implemented in Python’s Scipy package. Absolute integration tolerance is set to *atol* = 0.1*η*, where *η* is the migration strength. Initial conditions of species abundances are drawn from uniform distribution over [0, 1]. Perturbation strength is controlled using *ω* to satisfy ‖*ω* · *α*^(*d*)^‖_*F*_/‖*α*^(*r*)^‖_*F*_ = 0.05 throughout. Simulation parameters are summarized in Table I.

**Table I.** Simulation parameters used to create each of the figures. The parameters *γ*, *σ*_*K*_, *σ*_*m*_ are set to zero throughout.

| | $S$ | $M$ | $S/M$ | $\mu_c$ | $\sigma_c$ | $\mu_d$ | $\sigma_d$ | $K$ | $m$ | $\eta$ | $w$ (Sec. II D) |
| --- | --- | --- | --- | --- | --- | --- | --- | --- | --- | --- | --- |
| Fig. 1 (B) | 800 | 160 | 5 | 20 | 0.5 | 10 | 20 | 2 | 0.2 | $10^{-15}$ | - |
| Fig. 1 (C) | 800 | 160 | 5 | 20 | 4 | 10 | 20 | 2 | 0.2 | $10^{-15}$ | - |
| Fig. 2 | 1000 | 200 | 5 | 30 | 10 | 10 | 20 | 2 | 0.2 | - | - |
| Fig. 3 (A,B) | ∞ | ∞ | 5 | Varying | Varying | 10 | 20 | 2 | 0.2 | - | - |
| Fig. 3 (C) | Varying | Varying | 5 | 20 | Varying | 10 | 20 | 2 | 0.2 | $10^{-15}$ | - |
| Fig. 4 | 800 | 160 | 5 | 30 | Varying | - | - | 2 | 0.2 | $10^{-13}$ | $0.05 \cdot S$ |
| Fig. 5 | 800 | 160 | 5 | 50 | 20 | - | - | 5 | 0.2 | Varying | $0.05 \cdot S$ |
| Fig. 6 | 800 | 160 | 5 | 20 | 4 | 10 | 20 | 2 | 0.2 | $10^{-15}$ | - |
| Fig. 7 | 800 | 56 | 14.3 | 20 | 4 | 10 | 20 | 2 | 0.2 | $10^{-15}$ | - |
| Fig. 9 | 800 | 160 | 5 | 20 | 4 | 10 | 20 | 2 | 0.2 | 0 | - |
| Fig. 10 | Varying | Varying | 2 | 40 | 23 | 10 | 10 | 3 | 5 | $10^{-15}$ | - |
| Fig. 11 | ∞ | ∞ | 2 | 40 | Varying | 10 | 10 | 3 | 5 | $10^{-15}$ | - |

A Python code that run simulations of the ground model and its two variants, with example parameters for the Fixed Point and Persistent Dynamics phases, is given in: https://github.com/Itaydal/crm-chaos.

## Appendix C Cavity equations

To study the properties of a typical fixed point of the model, we use a variant of the cavity method [6, 9, 13, 27–29]. It proceeds by adding one new species and one new resource, along with newly sampled interactions between it and the rest of the system, creating an *S* + 1 species system with *M* + 1 resources. Then, by comparing the properties of a typical species of the old system with those of the newly added species and resource we get self-consistent equations for the macro-scopic variables *ϕ*, 〈*N*〉, 〈*N*^2^〉 where *ϕ* = *S*\**/S* is the fraction of living species, together with the properties of the resources.

Solving these self-consistent Eq. (C1) for range of parameters allows us to derive the phase diagram in Fig. In particular, the distinction between stable and non-equilibrium phases is done by solving for *ϕ* for some choice of control parameters, this determines the distribution of reduced interaction matrices, in Appendix D we calculate it’s stability. The transition into the unbounded growth phase is found at the divergence of (*N*).

### 1. Deriving species and resource distributions using cavity method

Introducing to the system new resource and species *R*_0_ and *N*_0_

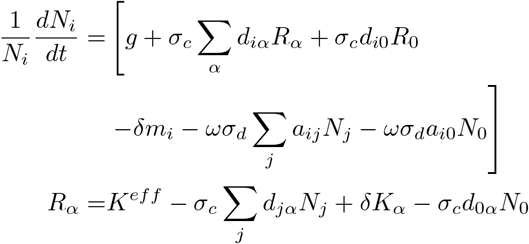

and the corresponding equations for *R*_0_ and *N*_0_ are

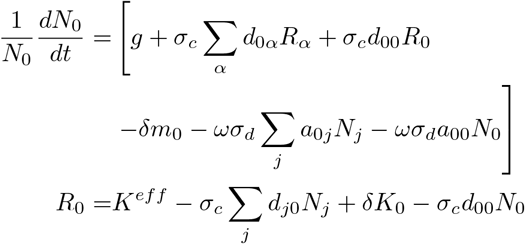

Denote the steady-state value of a quantity *X* by 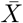, also denote by 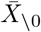 the steady-state value of *X* in the absence of the resource and species ‘0’.

Then we can define the following susceptibilities

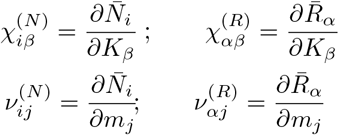

Since addition of single resource and species is a small (order *S*^−1^) perturbation we can write

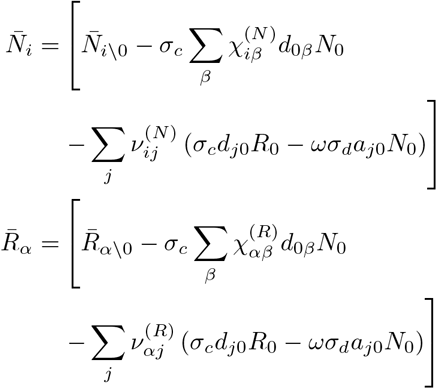

We can now plug in these expressions into the steady-state equations for *N*_0_ and *R*_0_. By taking leading order contributions to *S*^−1^, and take expectation value over expressions we get

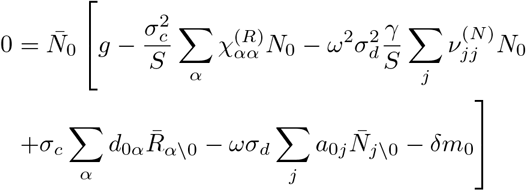

Notice that, to leading order in *S*^−1^, as sum of weakly interacting terms we can model the expression 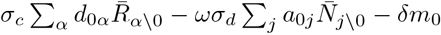 as a Gaussian random field with mean 0 and variance

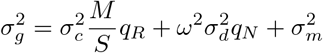

where

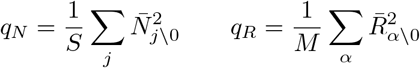

Let *z*_*N*_ be a Gaussian random field with zero mean and unit variance, and define the average susceptibilities

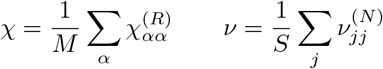

As there is no difference between species ‘0’ and the rest, we can emit the subscript ’0’ to and write the equation the fixed point abundance distribution

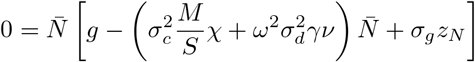

Following similar procedure for the resources yields

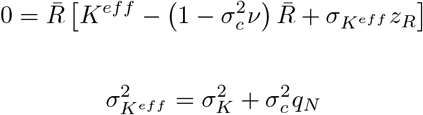

We can solve these equations and get

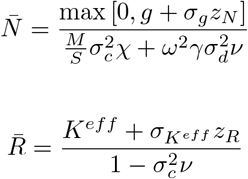

### 2. Self consistent equations

At this stage, our aim is to solve for {*ϕ*_*S*_, 〈*N*〉, 〈*R*〉, *q*_*N*_, *q*_*R*_, *χ*, *ν*} for a given set of control parameters *S, M, K, σ*_*K*_, *m*, *σ*_*m*_, *μ*_*c*_, *σ*_*c*_, *μ*_*d*_, *σ*_*d*_, *γ*, *ω*. To that end, it is helpful to define

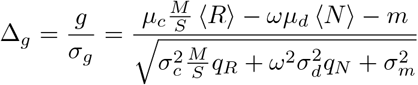

and the function

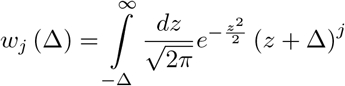

note that for 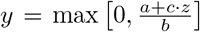 with *z* Gaussian random variable we have that

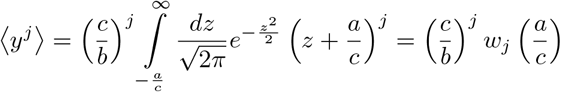

Taking the first two moments of the distributions 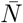 and 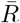, leads to the set of set consistent equations

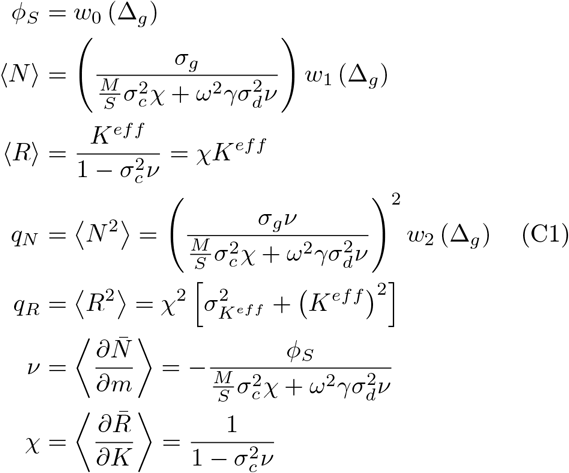

The expressions for *ν* and *χ* are derived by differentiating abundances distributions *N, R* with respect to *m* and *K* respectively and taking their expectation values.

To avoid singularities at the diverging phase (〈*N*〉 → ∞) we define 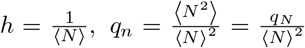. With these variables the self consistent equations read

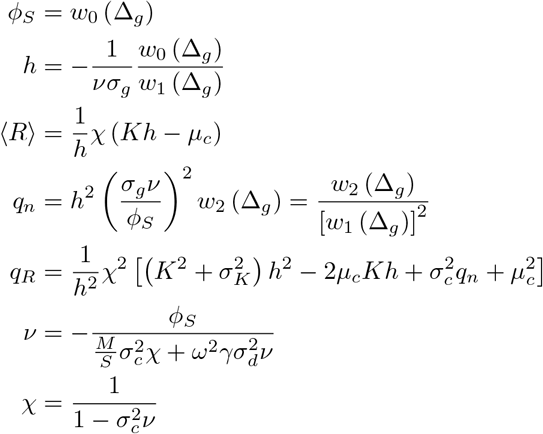

At this stage one has to find a self consistent solution for this set of equations. One possible approach would be to use a global numerical optimizer such as a basin-hopping algorithm to find a solution in the 7-dimensional parameter space spanned by *ϕ*_*S*_, *h, R, q*_*n*_, *q*_*R*_, ν, *χ*. This requires non-convex optimization in high dimension, which is not guaranteed to work. By some additional manipulation we were able to reduce it into a one dimensional non-convex optimization over the variable Δ_*g*_, as we now show.

Simplifying the expressions for the susceptibilities results with the third order polynomial for *ν* where the only unknown is *ϕ*_*S*_.

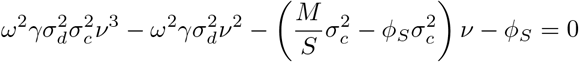

Note that *ϕ*_*S*_ only depends on Δ_*g*_, therefore one can span a grid of values for Δ_*g*_ and assigning the roots the above polynomial for each *ν*_*i*_ (Δ_*g*_) where *i* = 1, 2, 3. Plugging back into the expression for resources susceptibility leads to *χ*_*i*_ (Δ_*g*_).

Now, usingthe relations 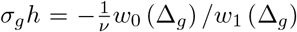 and 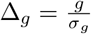 yields

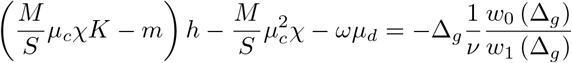

solving this for *h* (Δ_*g*_) leads to

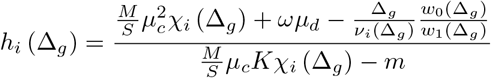

Rewriting the expression for Δ_*g*_ with the new variables *h, q*_*n*_

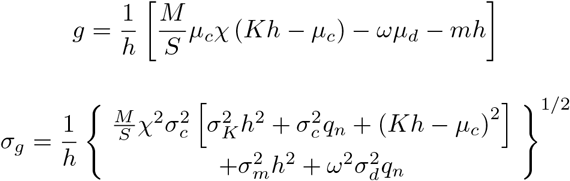

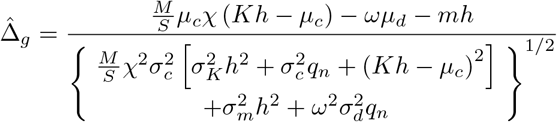

Finally, find values of Δ_*g*_ and *i* = 1, 2, 3 where 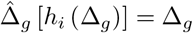. With these self consistent values for Δ_*g*_, *h*, ν, *χ* it is straight forward to then find *q*_*n*_, 〈*R*〉, *q*_*R*_.

## Appendix D Random Matrix Theory

Given the values of control parameters as described in Appendix A, the diversity *ϕ* = *S*\**/S* for the perturbed MCRM (Sec. II B) can be found as described in Appendix C. Here we define the random matrix ensemble corresponding to the reduced interaction matrix for given control parameters and diversity values. The main result of this appendix is the minimal eigenvalue real part of the ensemble Eq. (D1) in Appendix D 3. This in turn is used to distinguish between the stable and non-equilibrium phases in Fig. 3.

### 1. Random matrix theory and free probability

The linear stability of a fixed point is determined by the sign of the minimal eigenvalue of its interaction matrix. For randomly sampled interaction matrices, the problem of determining the sign of the minimal eigenvalue can be addressed with random matrix theory (RMT). A random matrix is a matrix whose elements are drawn from probability distribution, known as an ensemble. One of the main uses of RMT is to determine what the spectrum of a typical matrix drawn from such ensemble would look like, and in particular its minimal eigenvalue. Below we describe the key steps taken to find t he m inimal e igenvalue o f t he p articular ensemble at hand. For a detailed review of these techniques see [30].

A central object in RMT is the Green function of an ensemble, also known as a Resolvent or Stieltjes transform. For an *N × N* random matrix *H*, the Green function is defined as

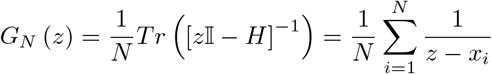

where *x*_1_, … *x*_*N*_ are the eigenvalues of *H*. Since *H* is a random matrix, *G*_*N*_ (*z*) is a random complex function with poles at locations *x*_*i*_. There are several methods for deriving the green function for a given ensemble, for details see [30]. Averaging over *H* and taking the thermodynamic limit (*N* → ∞),

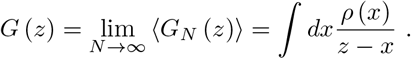

At the thermodynamic limit the set of eigenvalues *x*_1_, … *x*_*N*_ becomes the eigenvalue density *ρ* (*x*) for the ensemble. Using the Sokhotski-Plemelj formula one can extract the eigenvalue density *ρ* (*x*) from the green function *G*(*z*) as follows

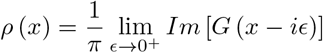

In this work, we want to calculate properties of sums of random matrices (the sum of the MCRM interactions and direct interactions). In general, random matrices do not commute, and the spectrum of the sum matrix isn’t simply the sum of the spectra. Therefore it is hard to calculate the spectrum of random matrices sum even given access to the Green functions of the ensembles. Free probability is a tool generalizing the notion of random variable independence to the field of random matrices. Analogous to statistical independence for random variables, two random matrix ensembles may exhibit the ‘freeness’ property, the precise definition can be found in [30].

Free probability provides us with a prescription for deriving the Green function of the ensemble sum given the Green function of the summed ensembles exhibiting the freeness property. This is analogous to the convolution law for random variable sum. It proceeds as follows. First, define the complex valued blue function to be the functional inverse of the green function

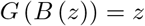

Now, given the blue function of the two ensembles *B*_1_ (*z*) *, B*_2_ (*z*), the blue function of the sum ensemble reads

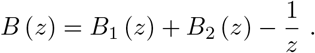

Finally, to find the green function of the sum ensemble, invert the blue function above using the relation

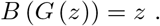

### 2. Wishart, GOE and Ginibre ensembles

The perturbed MCRM interaction matrix appearing in Sec. II C consists of a sum of two matrices:

1. Resource competition interaction matrix - Wishart ensemble

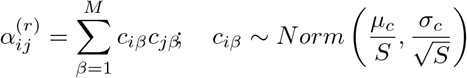

with the blue function

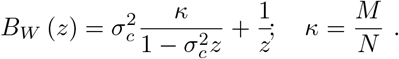
2. Direct competition interaction matrix - Ginibre ensemble

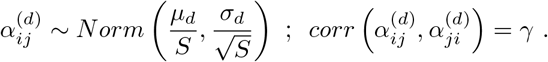

In general (for *γ* ≠ 1) a matrix drawn from the Ginibre ensemble is not Hermitian and therefore has a complex valued spectrum. Non Hermitian ensembles call for a generalization of the Green function. Concretely, in these cases the Green function would be a Quaternionic valued function leading to much more complicated calculations. Luckily, the Ginibre ensemble can be assembled as the sum of two independently distributed matrices from the Gaussian orthogonal ensemble (GOE) with complex prefactors [31]. This representation of the Ginibre ensemble allows for great simplification following method by [32].

Given two matrices *H, H′*, with elements drawn independently from *H*_*ij*_ ~ *Norm* (0, *σ*^2^*/N*) and symmetrize (*H* + *H*^*T*^) /2. The Ginibre matrix *α*^(*d*)^ can be written as

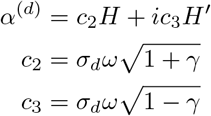

where *ω* is the aggression factor maintaining a constant direct perturbation strength as described in II B. The blue function of the GOE with real value prefactor *c* is given by

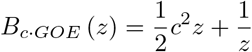

Finally, the ensemble for the perturbed interaction matrix *α* = *α*^(*r*)^ + *ω* · *α*^(*d*)^ can be written as

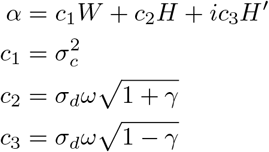

with the blue functions for the real and imaginary parts

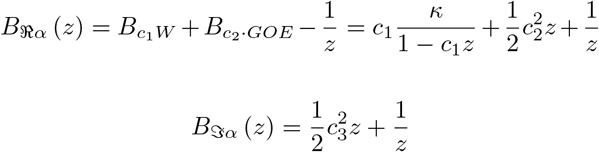

### 3. Calculating the minimal eigenvalue of the matrix sum

In this section we derive the minimal eigenvalue real part for the ensemble describing the reduced interaction matrix of the perturbed MCRM in Sec. II B. This is the main result of this appendix, then being utilized to find the phase diagram in Fig. 3.

In this section we follow the method by [32] to find the spectrum of the sum of Hermitian random matrices with imagery prefactors. Using this method one can derive the spectrum of a non-Hermitian random matrix *H*_1_ + *iH*_2_ comprised of two Hermitian matrices *H*_*1*_, *H*_2_, without having to go through cumbersome calculations green functions quaternionic.

Still, it is hard to find the entire spectrum of this ensemble for *α*. We simplify the problem further, trying to find just the minimal real part of the complex spectrum, given as a particular case of the equations for spectrum support contour on the complex plane.

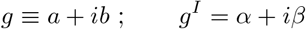

By [32], Eq. (77),

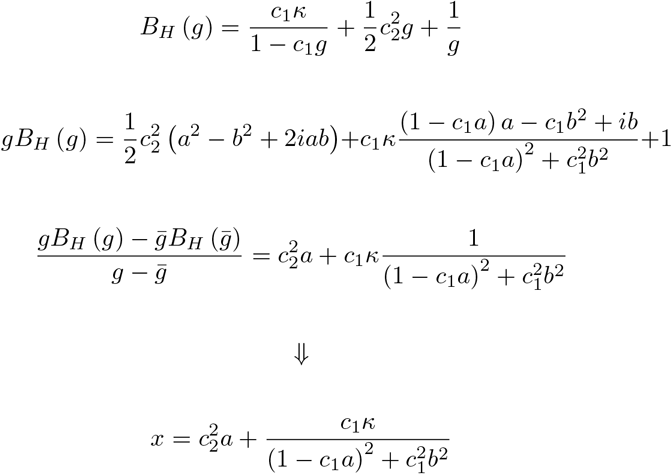

By [32], Eq. (78),

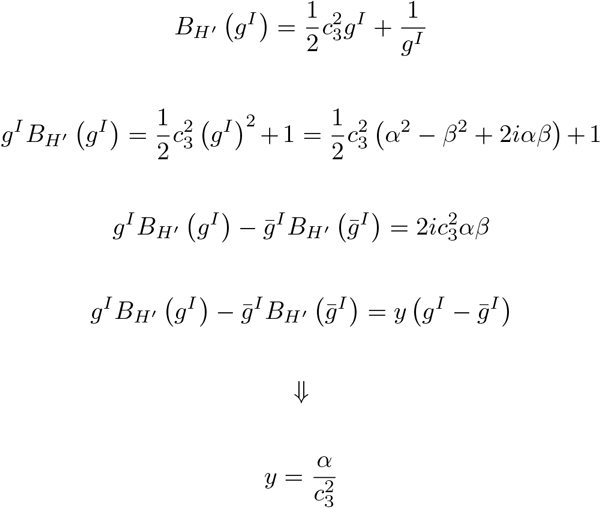

And from [32], Eq. (74),

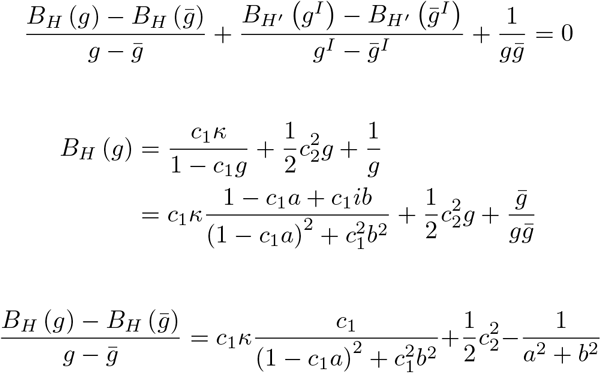

Using [32], Eq. (63) we have 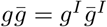

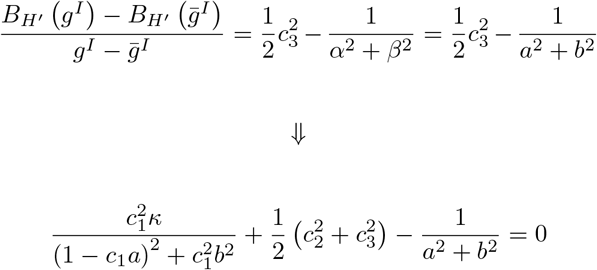

Combining [32], Eqs. (63,74,77) we get the set of coupled equations

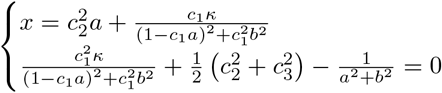

According to [32], Eq. (93) the spectrum contour equation is given by 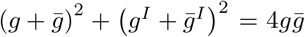. Now, fo-cusing on the real part of the contour, given by *y* = 0 combined with [32], Eq. (78) leads to *α* = 0. Therefore the contour equation reduce to

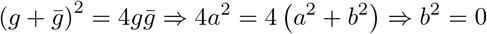

Plugging that back to the set of coupled equations above, yields the polynomial equation for *a*

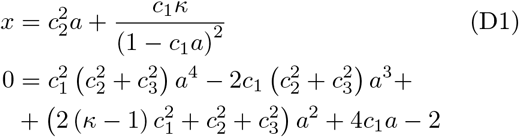

Substituting back the real *a* roots into Eq. (D1) to get the minimal and maximal eigenvalue real parts of the ensemble. By doing so determining the linear stability of the perturbed MCRM model at Eq. (2).

## Appendix E Species abundance distribution

At fixed points of resource-consumer models, species diversity is limited by the number of limiting resources (*S*\* ≤ *M*), according to the competitive exclusion principle [25]. In contrast, non-equilibrium states are not bound by the exclusion principle and can exceed this limit, i.e. *M < S*\*. In this section we show that for the system described in Eq. (3) this is indeed the case, even if the migration is very small (in the *η* → 0 limit).

As discussed in Sec. II E, simulations with migration *η*, show a typical species abundance probability distribution, see Fig. 5(A), with a power law as the abundance distribution between an upper value *N*_*u*_, and lower value determined by the migration floor *η*. We write this probability distribution as

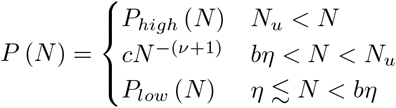

The power law behavior is parameterized by *c, ν* that may dependent on the migration *η*. The lower part *P*_*low*_(*N*) is the abundance distribution for species that are maintained thanks to migration. We define this region to go up to *bη* with a (somewhat arbitrary) constant value *b*.

Our interest is in when and how species diversity goes beyond the competitive exclusion bound in this probability distribution. Let us define *C*_*high*_, *N*_*CE*_, *C*_*CE*_ as

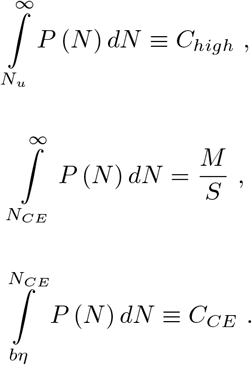

Note that if competitive exclusion holds and *η* = 0, there are no species in the range 0 *< N < N*_*CE*_. That is, species are either extinct and concentrated at *N* = 0 of *P* (*N*), or have an abundance above *N*_*CE*_. Therefore, to demonstrate that the species diversity can exceed the competitive exclusion limit in chaotic states we show that 0 *< C*_*CE*_ at the limit of vanishing migration.

For simplicity of the analysis we replace *P*_*low*_ by extending the power law and introducing a new lower cutoff at *aη*, to preserve the area under the curve of this lower region. We still treat abundances below *bη* as species maintained solely by migration. The simplified probability distribution reads

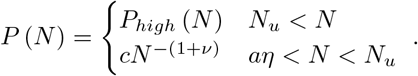

From normalization of *P* (*N*) we solve for 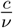 to find

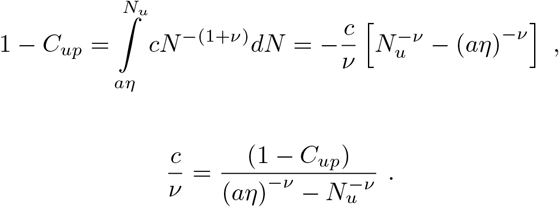

Next, we express *N*_*CE*_ as

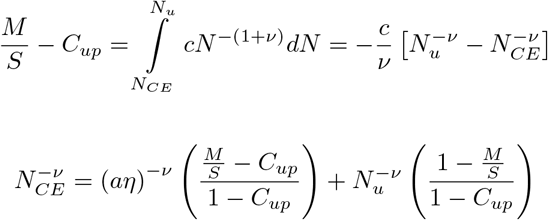

Finally, *C*_*CE*_ takes the form

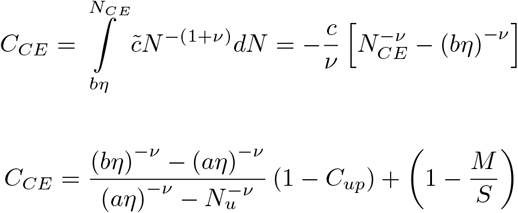

At the limit *η* → 0

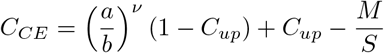

This equation expresses *C*_*CE*_ as a function of the parameters of the probability distribution (*a, b, ν, C*_*up*_), and the number of species *M* and resources *S*. Simulations shows that 0 ≤ *ν* ≪ 1 (possibly vanishing) at the limit *η* → 0. Assuming that *M/S <* 1 (there are more species in the pool than resources), we conclude that 0 *< C*_*CE*_, hence non-equilibrium states of consumer-resource models can sustain diversity exceeding the competitive exclusion limit.

Finally, a note regarding the diversity, compared with the diversity as predicted by the cavity solution described in Appendix C. That solution is only exact when the system reaches a unique stable equilibrium. Elsewhere it is an approximation; from simulations of the first model variant in Sec. II B, we find that the cavity solution is lower than the one described in this Section, see Fig. 6.

**Figure 6.**
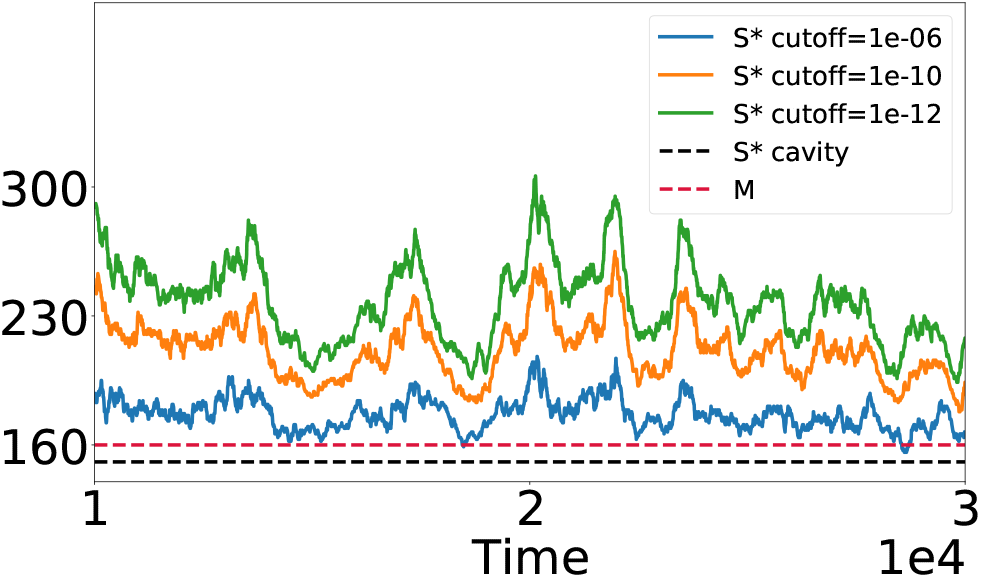
Number of coexisting species *S*\*, as a function of time in the persistent dynamics phase. A species is counted in the standing diversity *S*\* if its abundance is above the level given in the legend. As can be seen, the number coexisting species exceeded the amount of resources for large range of abundance levels.

**Figure 7.**
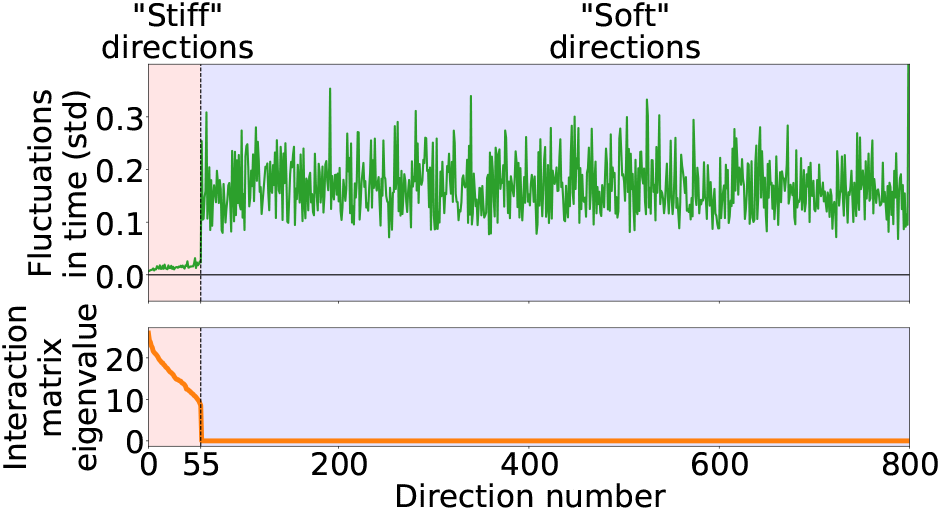
(Top) Fluctuations over time along the eigenvector directions of the resource interaction matrix *α*^(*r*)^. (Bottom) Corresponding eigenvalues *λ*_*i*_ of the spectral decomposition of *α*^(*r*)^. There is a clear distinction between “Stiff” directions showing little fluctuations c orresponding to fi nite positive eigenvalues, and “Soft”, strongly fluctuating marginal directions with corresponding zero eigenvalues.

## Appendix F Stiff and soft fluctuation directions

It is interesting to see whether the fluctuations of the abundances are directly related to the marginal directions of the MCRM fixed points. To do that, we rotate the vector 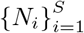 in a way that will separate the “stiff” degrees of free*i*d=o1m, lying in the non-marginal directions of a MCRM fixed point, and the “ soft” degrees of freedom at the marginal dimensions. This is done by rotating with an orthogonal matrix *O* the abundance vector 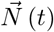 from a simulation of the perturbed MCRM in it’s chaotic phase. The orthogonal matrix *O* is obtained from the spectral decomposition of the unper-turbed interaction matrix 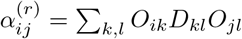 where *D* is diagonal matrix. Note that since *α*^(*r*)^ is a symmetric positive-semi-definite matrix, its eigenvalues 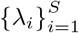 are real valued and non-negative.

Denote by 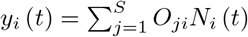 the rotated degrees of freedom. These are a combination of species abundances at time *t*. Define the fluctuation over time in direction *i* to be std 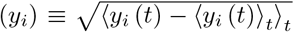, where 〈*…*〉_*t*_ denotes time average over a Δ*t* = 10^4^ interval. Plotting std (*y*_*i*_) sorted by the eigenvalue *λ*_*i*_, shows that the fluctuations in the “soft” directions (where *λ*_*i*_ = 0) are consistently and significantly larger than in the “stiff” directions (where *λ*_*i*_ > 0), see Fig. 7.

## Appendix G Isolated systems (no migration)

Here we discuss cases where there is no migration from an external “mainland” pool of species. We consider both a single community, and a a meta-community, a setting in which multiple well-mixed communities are coupled by migration. We show that meta-communities can allow for persistent dynamics over long times, even in the absence of external migration, and for finite population sizes. In isolated well-mixed communities, simulations show that extinctions drive the system to a fixed point, with diversity a little below the competitive exclusion bound.

The dynamics of the meta-community are a set of differential equations for 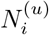 describing the abundance of the *i*-th species in the *u*-th community,

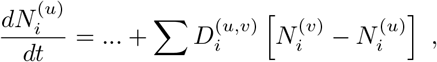

where the “…” refers to the terms in the RHS of Eq. (2) applied to 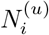, with *η*_*i*_ = 0. A species is considered extinct and removed from the system when its abundance 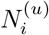 goes below some cut-off *N*_*c*_ in all communities *u*, corresponding to the inverse of the population size.

Fig. 8 shows the dynamics at late times of a few representative abundances 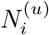, showing persistent fluctuations in a meta-community comprised of 8 communities with *S* = 400 species and *M* = 80 resources. The model in each patch corresponds to that in Sec. II B. The resource interaction matrix *α*^(*r*)^ has *μ*_*c*_ = 30 and *σ*_*c*_ = 6. The matrix *α* is very similar but not identical between the different communities, with correlation *ρ* = 0.997 between the 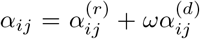 for the same *i, j* in different communities. Direct interaction matrix *α*^(*d*)^ is drawn independently for each community with *μ_d_* = 10, *σ_d_* = 20 and *γ* = 0. As in the main text, *ω* is determined to satisfy perturbation strength of 0.05. Coupling between communities set to be *D* = 10^−4^. Cutoff abundance is taken to be *N*_*c*_ = 10^−20^.

A simulation of a single, well-mixed community is shown in Fig. 9, along with the diversity as a function of time. The diversity drops, until the system reaches a fixed point with *S*\* a little below *M*, almost saturating the competitive exclusion bound. Simulation parameters are specified in Tab. I and are similar to that in Fig. 1(C) but with *η* = 0.

**Figure 8.**
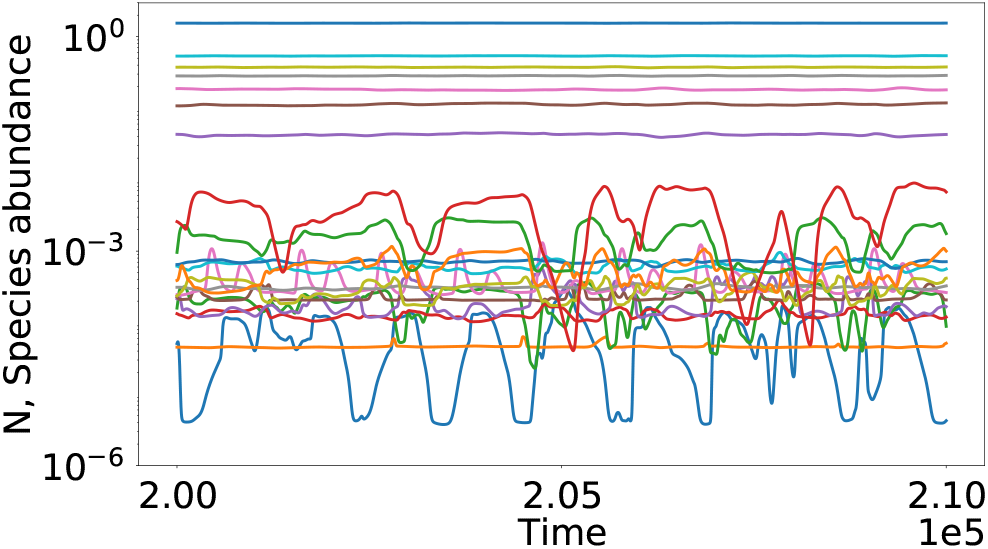
Dynamics of a meta-community composed of 8 communities coupled by migration, at late times. Persistent abundance fluctuations are shown, which do not go below some value, showing that even finite populations can persist for very long times. 20 representative species are plotted.

**Figure 9.**
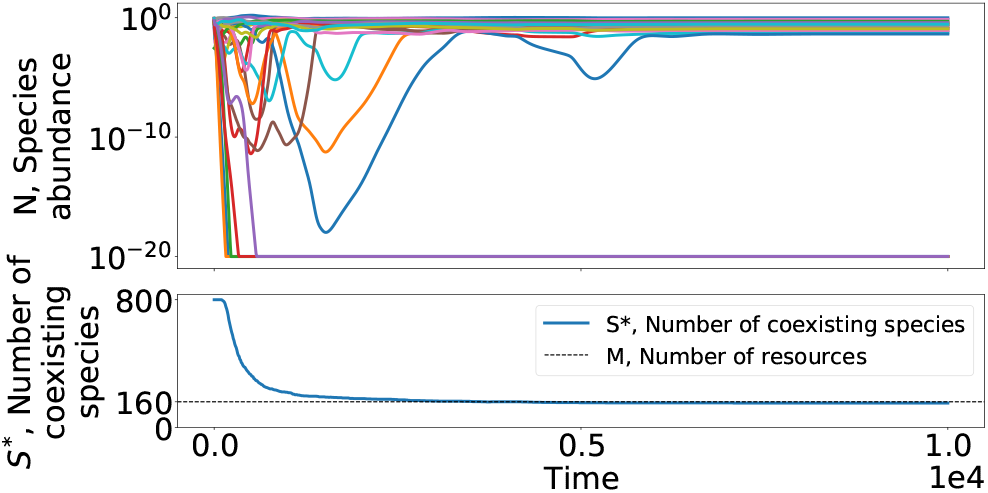
Dynamics of a single community without external migration at the chaotic phase. Species abundances initially fluctuate, and some go extinct. A fixed point is reached once the diversity goes a little below the number of resources *M*.

## Appendix H Symmetric additional interactions

Here we consider a setting similar to that in Sec. II B, where additional interactions 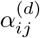 are added the ground model. The difference is that here they are taken to be symmetric, 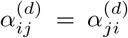. This difference is important, since in this case, the entire interaction matrix *α* is symmetric. This means that the dynamics admit a Lyapunov function, and always reach a fixed point. A similar situation, with symmetric random Lotka-Volterra interactions (in this work’s terminology, *α* = *α*^(*d*)^) has been studied in [12]. There, a fixed point phase was found. Beyond it lies a *critical* phase, characterized by many alternative equilibria, all of them close to marginal stability, namely such that the minimal eigenvalue *λ*_*min*_ → 0 as *S* → ∞. Specifically, it was found that *λ*_*min*_ ∝ *S*^−2/3^.

Here we find precisely the same phenomenology, with a fixed point phase. Beyond it simulations show that the system possesses multiple alternative equilibria. Furthermore, the minimal eigenvalue was measured for multiple values of *σ*_*c*_ and *S*, and averaged over many runs.

**Figure 10.**
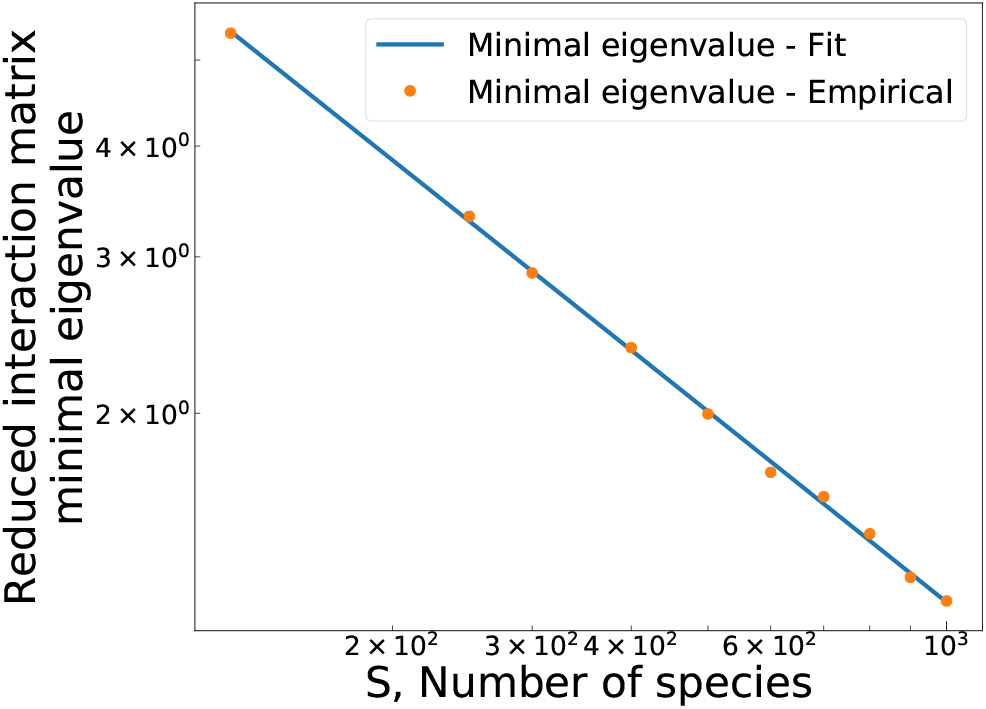
Fitting the minimal eigenvalue of the reduced interaction matrix to *λ*_*min*_ (*S*) = *a · S*^*b*^ + *λ*_∞_. Parameters as in Fig. 11, with *σ*_c_ = 23.

**Figure 11.**
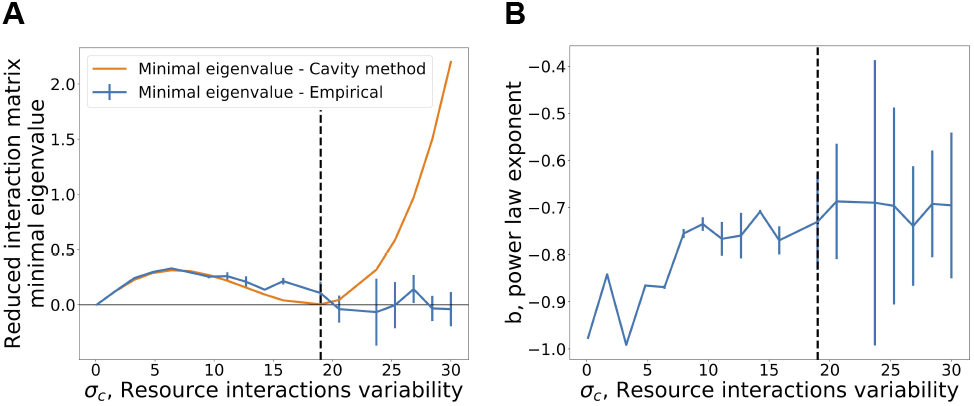
Minimal eigenvalue of the reduced interaction matrix with symmetric direct interactions perturbation (*γ* = 1). Minimal eigenvalue at *S → ∞* is obtained from a fit to *λ*_*min*_ (*S*) = *a · S*^*b*^ + *λ*_∞_, see Fig. 10. Shown are (A) The minimal eigenvalue *λ*_∞_, and (B) the power law exponent *b*.

For each value of *σ*_*c*_ it was fit to *λ*_*min*_ (*S*) = *a* · *S*^*b*^ + *λ*_∞_, where *a, b, λ*_∞_ depend on *σ*_*c*_, see Fig. 10. The results for *b, λ*_∞_ are shown in Fig. 11. We find that beyond the unique fixed point phase the results are very different from the simple cavity solution for this case, and consistent with *λ*_∞_ = 0 and *b* = −2/3 which was predicted for the random Lotka-Volterra setting.

